# Quantitation of brain tumour microstructure response to Temozolomide therapy using non-invasive VERDICT MRI

**DOI:** 10.1101/182675

**Authors:** Tom A. Roberts, Harpreet Hyare, Giulia Agliardi, Ben Hipwell, Angela d’Esposito, Andrada Ianus, James O. Breen-Norris, Rajiv Ramasawmy, Valerie Taylor, David Atkinson, Shonit Punwani, Mark F. Lythgoe, Bernard Siow, Sebastian Brandner, Jeremy Rees, Eleftheria Panagiotaki, Daniel C. Alexander, Simon Walker-Samuel

## Abstract

There has been slow progress in the development of new therapeutic strategies for treating brain tumours, partly because assessment of treatment response is difficult and largely reliant on simple bi-dimensional measurements of MRI contrast-enhancing regions. Hence, there is a clinical need to develop improved imaging techniques for monitoring treatment response. In this study, we evaluate VERDICT (Vascular, Extracellular and Restricted Diffusion for Cytometry in Tumors) MRI in mouse glioblastomas for the quantification of tumour microstructure and assessment of response to Temozolomide (TMZ) chemotherapy, and, we investigate the feasibility of applying VERDICT MRI in a range of human gliomas. VERDICT MRI detected response to TMZ earlier than structural and apparent diffusion coefficient (ADC) measurements. A significant reduction in the cell radius parameter was detected three days earlier than ADC and six days earlier than structural MRI. Histological analysis showed the same trend as VERDICT of decreased intracellular volume fraction in the TMZ-treated mice. Vascular volume fraction was not altered by TMZ, which was consistent with optical projection tomography measurements. In patients, glioblastoma compartmental volume fractions showed good agreement with mouse glioblastoma parameters. The VERDICT parameters varied across the human gliomas, with raised intracellular volume fraction in the oligodendrogliomas and elevated cell radius in both low-grade tumours subtypes. In conclusion, our results suggest that VERDICT MRI is more sensitive at detecting TMZ response than structural or ADC measurements. In patients, VERDICT is feasible within clinical scan times, and performed best at characterising glioblastoma. Further optimisation should improve assessment of different glioma subtypes.

## INTRODUCTION

Gliomas are the most common type of primary brain tumour in adults with an annual incidence of 4-5/100,000 people. For newly diagnosed glioblastoma (GBM), the most common and malignant of the gliomas, no treatment has yet been shown to be more effective than maximal safe surgical resection followed by chemoradiation and adjuvant chemotherapy with Temozolomide (TMZ) (1). There has been only a modest improvement in outcome over the last 10 years with three year survival increasing from about 4% in 1999-2000 to 10% in 2009-2010 (2).

Despite new therapeutic strategies inhibiting angiogenesis and more recently immunotherapy (3), progress has been limited by the inherent difficulties in assessing treatment response in clinical trials. Traditional imaging of response utilizes measurements of enhancing tumour on T1-weighted MRI (4) and these have been confounded by the phenomena of *pseudoprogessi*on, whereby an increase in tumour volume, oedema and enhancement, shortly after completion of chemoradiation treatment is indistinguishable from progressive tumour (5,6) and *pseudoresponse*, whereby a dramatic reduction in tumour enhancement following treatment with anti-angiogenic agents is thought to be due to vascular normalization rather than a true anti-tumour effect (7,8).

Although the current most widely used response assessment criteria for high-grade gliomas known as the RANO criteria (Response Assessment in Neuro-oncology) corrects for a number of these complexities (9), there is still reliance on bi-dimensional measurements for contrast enhancing tumours. As enhancement in high grade gliomas reflects only one aspect of the biology of these tumours, i.e. blood brain barrier breakdown (10), there is an urgent need to develop quantitative and robust imaging technologies to better assess tumour burden and response to treatment.

Diffusion weighted imaging (DWI), which is the technology that underpins VERDICT, has been widely shown to offer additional utility in brain tumours beyond tumour volume imaging (11). DWI is sensitive to the restriction of water diffusion by tissue microstructure, and can therefore be used to probe structures below the resolution of the image acquisition. It does not require the administration of a contrast agent and does not require additional specialist equipment. The apparent diffusion coefficient (ADC) in gliomas has been shown to inversely correlate with tumour cell density (12,13), likely to be due to restriction of water motion in tightly packed cells, making DWI a technique of great interest for both grading and detecting disease recurrence (14) and for differentiating pseudoprogression from progressive disease (15,16).

DWI data are usually quantified by fitting a single exponential function to images acquired with increasing diffusion weighting, to estimate the ADC (17). More sophisticated models have been proposed, which include, but are not limited to: intravoxel incoherent motion (IVIM) (18), diffusion tensor imaging (DTI) (19), diffusion kurtosis imaging (20) and neurite orientation dispersion and density imaging (NODDI) (21).

More recently, VERDICT has shown promise for non-invasively quantifying microstructural changes in tumours. To date, its use has been investigated in colorectal cancer (22) and prostate cancer (23), and has shown a good ability to distinguish tumour from healthy tissue and to accurately quantify cell size and density. VERDICT utilises multiple MRI diffusion gradients to probe the microscopic motion of water molecules tumours at a range of length scales, and a three-compartment biophysical model is fitted to the DWI signal, providing estimates of intracellular volume fraction (*f_ic_*), extracellular-extravascular volume fraction (*f_ees_*), intravascular volume fraction (*f_v_*) and cell radius (Figure S1). These parameters are particularly relevant to cancer, where cell proliferation is elevated and angiogenesis is a key factor in tumour progression.

To date, the ability of VERDICT MRI to evaluate brain tumours has not been investigated. In this study, we have investigated the use of VERDICT in a GL261 GBM mouse model (24) to assess pre-therapy microstructure and to measure response to Temozolomide therapy. We have then applied the optimised VERDICT model to a cohort of human adult gliomas in an initial feasibility study. Our hypothesis was that VERDICT MRI could detect tumour microstructural changes in response to therapy before macroscopic changes, with potential as a therapeutic marker in human glioma treatment trials.

## MATERIALS AND METHODS

### Mice and glioma cell implantation

All *in vivo* experiments were performed in accordance with the UK Home Office Animals Scientific Procedures Act, 1986 and United Kingdom Coordinating Committee on Cancer Research (UKCCCR) guidelines (25).

Female, 8-weeks old, C57BL/6 mice were injected with 2x10^4^ GL261 mouse glioma cells. Mice were anesthetized with 4% isoflurane in an induction box and then transferred to a stereotactic frame (David Kopf Instrument, Tujunga, CA), where anaesthesia was delivered through a nose cone and maintained at 2%. The head was sterilised with 4% chlorhexidine and the skin was cut with a sterile scalpel to expose the skull. Coordinates were taken using a blunt syringe (Hamilton, 75N, 26s/2”/3, 5 μL): 2mm right and 1mm anterior to the bregma, corresponding to the right caudate nucleus. A burr hole was made using a 25-gauge needle. The Hamilton syringe was lowered 4mm below the dura surface and then retracted by 1mm to form a small reservoir. 2x10^4^ GL261 cells were injected in a volume of 2 μL over two minutes. After leaving the needle in place for 2 minutes, it was retracted at 1 mm/min. The burr hole was closed with bone wax (Aesculap, Braun) and the scalp wound was closed using Vicryl Ethicon 6/0 suture.

### Study design and administration of Temozolomide

Mice were implanted with glioma cells for imaging. At 13-days post inoculation (day 0), mice were randomly assigned to control or TMZ-treated groups (n = 12 for each), and baseline imaging was carried out. Immediately after scanning, and following recovery from anaesthesia, mice in the TMZ group were administered with a first dose of Temozolomide (Temodar, MerckKenilworth, NJ) by oral gavage (130 mg/kg in vegetable oil). Two further doses were given on consecutive days, to a total dose of 490 mg/kg. Mice in the control group received sham doses of vegetable oil according to an equivalent regimen. MRI was carried out every three days, to a final timepoint at 22-days post tumour injection (day 9).

### MRI acquisition

Mouse MRI measurements were performed on a 9.4T horizontal bore scanner (Agilent Technologies, Santa Clara, CA) with a 205/120/HD 600 mT/m gradient insert. RF transmission was performed with a 72 mm inner diameter volume coil and a 2-channel receiver coil (RAPID biomed, Ripmar, Germany). Mice were anaesthetised with 1.5-2.0% isoflurane in 2 l/min oxygen, and positioned prone in a cradle for imaging. The head was positioned within an MR-compatible head-holder and secured with plastic ear bars to minimise motion artefacts. Body temperature was maintained at physiological temperature using a hot water system and monitored using a rectal probe (SA Instruments, Stony Brook, NY). Respiration was monitored using a neonatal apnoea pad taped to the abdomen of the animal. An intraperitoneal infusion line was used for administration of gadolinium contrast agent (Magnevist, Bayer, Leverkusen, Germany).

### Imaging Protocol

Following shimming, tumours were localised using a structural T2-weighted spin-echo sequence. For VERDICT, diffusion-weighted images were acquired in a coronal orientation using a 3-shot spin-echo echo planar imaging (EPI) sequence, which included the following parameters: TR = 3s, TE = min, data matrix = 64x64, FOV = 20x20mm, shots = 3, slice thickness = 0.5mm, slices = 5, averages = 2. In total, 46 diffusion weightings (each of 3 directions) were acquired in addition to a 42 direction DTI acquisition (b = 1000 s/mm^2^). Specific gradient combinations are detailed in Table 1. TE was minimised for all scans to maximise signal-to-noise. To correct for signal changes caused by this variation in TE, an accompanying b ~ 0 s/mm^2^ (B_0_) image was acquired for every combination of diffusion gradients. Total imaging time for VERDICT was 70 minutes. Following VERDICT, mice were injected with 0.6 mmol/kg of gadolinium-DTPA. After 10 minutes to allow for the contrast agent to circulate, slice-matched T1-weighted spin echo EPI images were acquired. Tumour regions of interest (ROI) were drawn based on these images and used during the quantification of the diffusion data.

### Quantification of DWI data

VERDICT models the water signal from three non-interacting tissue compartments (22). In brief, a BallSphereStick model (Figure S1) characterises the diffusion signal, in which S_sphere_ is the signal from restricted water (intracellular), S_stick_ is the signal from blood (intravascular) and S_ball_ is the signal from the extracellular-extravascular space. The total signal within a voxel is the weighted sum of the contributions from each compartment, according to:

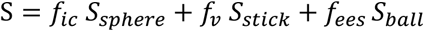

Further details on the functional form for each component can be found in Panagiotaki *et al* (22).

Model-fitting was carried out in MATLAB (Mathworks, USA) using the open-source Camino toolbox (26). In total, four parameters were estimated in this study: *f_ic_, f_ees_, R* (cell radius) and *dv* (intravascular diffusivity). For model stability, the intracellular and extracellular-extravascular diffusivities were fixed: *d_ic_* = 1x10^-9^, *d_ees_* = 2x10^-9^ m^2^/s respectively. The intravascular volume fraction was not fitted and instead was calculated as *f_v_* = 1 − (*f_ic_* + *f_ees_*).

Whole-brain ADC and VERDICT parameter maps were generated for visualisation, however, as the VERDICT model is an unsuitable descriptor of normal brain tissue, only parameter values within tumour ROIs were used in further analysis. In glioblastoma patients, ROIs were drawn around three separate regions corresponding to the tumour core, tumour rim and peri-tumoural zone. Analyses were performed independently on each of these regions.

### Optical Projection Tomography

Optical projection tomography (OPT) of complete brains was used to quantify the blood volume of tumour tissue, in a subset of mice (n = 1 control, n = 1 TMZ-treated), for comparison with *in vivo* MRI data. The full method for fluorescently-labelling the vasculature, clearing the mouse brains, imaging, and image processing is detailed in the Supplementary Material.

### Histology

Intracellular volume fraction was quantified from histology for comparison with VERDICT MRI using an in-house MATLAB script. After the final MRI scan, mouse brains were extracted (n = 8 control, n = 6 TMZ-treated), immersion fixed in 4% PFA, and then sliced in a coronal orientation and stained with hematoxylin and eosin (H&E). A k-means threshold was applied to the H&E slices to estimate the ratio of the stained brain tumour cells to unstained extracellular-extravascular space, for comparison with the VERDICT *f_ic_* parameter.

### Patients and MRI acquisition

The human MRI component of the study was performed with local ethics committee approval and patients provided written informed consent. Nine untreated patients with brain tumours (3 glioblastoma (WHO grade IV), 3 astrocytoma (WHO grade II/III) and 3 oligodendroglioma (WHO grade II)) were scanned at 3T (Achieva, Philips) using an 8-channel head coil. Tumours were localised using T1- and T2-weighted structural MRI sequences. For VERDICT MRI in patients, nine diffusion weightings were acquired (3-orthogonal directions, b = 80-3000 s/mm^2^) in a protocol that minimised scan time whilst maximising the range of diffusion times covered. A single-shot spin echo EPI readout was used with the following parameters: TR = 3.7s, TE = min, FA = 90°, DM = 92^2^, voxel size = 2.5mm^3^, slices = 33, averages = 1. A reduced 15-direction DTI scan was also acquired (b = 700 s/mm^2^). Specific diffusion gradient combinations are detailed in Table 2. Total acquisition time was 12 minutes. DWIs were normalised to b = 0 s/mm^2^ (B_0_) images acquired with the same TE. Five patients immediately underwent a second set of same-session scans for assessment of repeatability. Prior to VERDICT analysis, diffusion images were registered and corrected for eddy current effects using the ECMOCO toolbox (27) in SPM (28). Tumour masks were created by manual segmentation of B_0_ images. For the GBM tumours, additional ROIs were created for the necrotic tumour core (GBM Core), enhancing rim (GBM Rim) and GBM perilesional T2-weighted signal abnormality (GBM Peri).

### Biopsy

Biopsies were retrieved from seven of the brain tumour patients. Image processing and analysis was performed on H&E biopsy samples to estimate nuclei volume fraction for comparison with VERDICT estimates of intracellular volume fraction. Nucleus volume fraction was estimated, rather than cellular volume fraction, due to the difficulty in delineating cell boundaries on biopsy samples. In principle, nucleus fractional volume is a valid marker of intracellular volume fraction. The full biopsy image processing pipeline is detailed in the Supplementary Material.

### Statistics

For the majority of comparisons in the results, Mann-Whitney-U statistical tests were performed to assess significance. For repeatability measurements of VERDICT parameters in patients, significance was assessed using a Wilcoxon matched-pairs signed rank test. The repeatability coefficient (*RC*) was also calculated for each parameter, which represents the 95% confidence interval of the difference in the two trials. RC is given by:

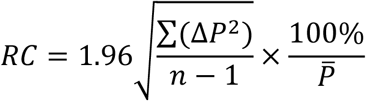

where *ΔP* is the change in parameter value between trials and 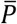 is the mean estimate of the VERDICT parameter between trials. For interpretation, the *RC* provides an indication of the change required to observe a difference above variation.

## RESULTS

### GL261 mouse glioma response to Temozolomide

GL261 GBM tumours were conspicuous in diffusion weighted images (b = 1000 s/mm^2^) where the tumour had a lower signal than the rest of the brain (Figure 1). Tumours were also visible on T1-weighted post-gadolinium images, and were characterised by increased signal within the tumour region, compared to normal brain, reflecting raised blood vessel density, blood flow and/or vessel permeability.

**Figure 1:**
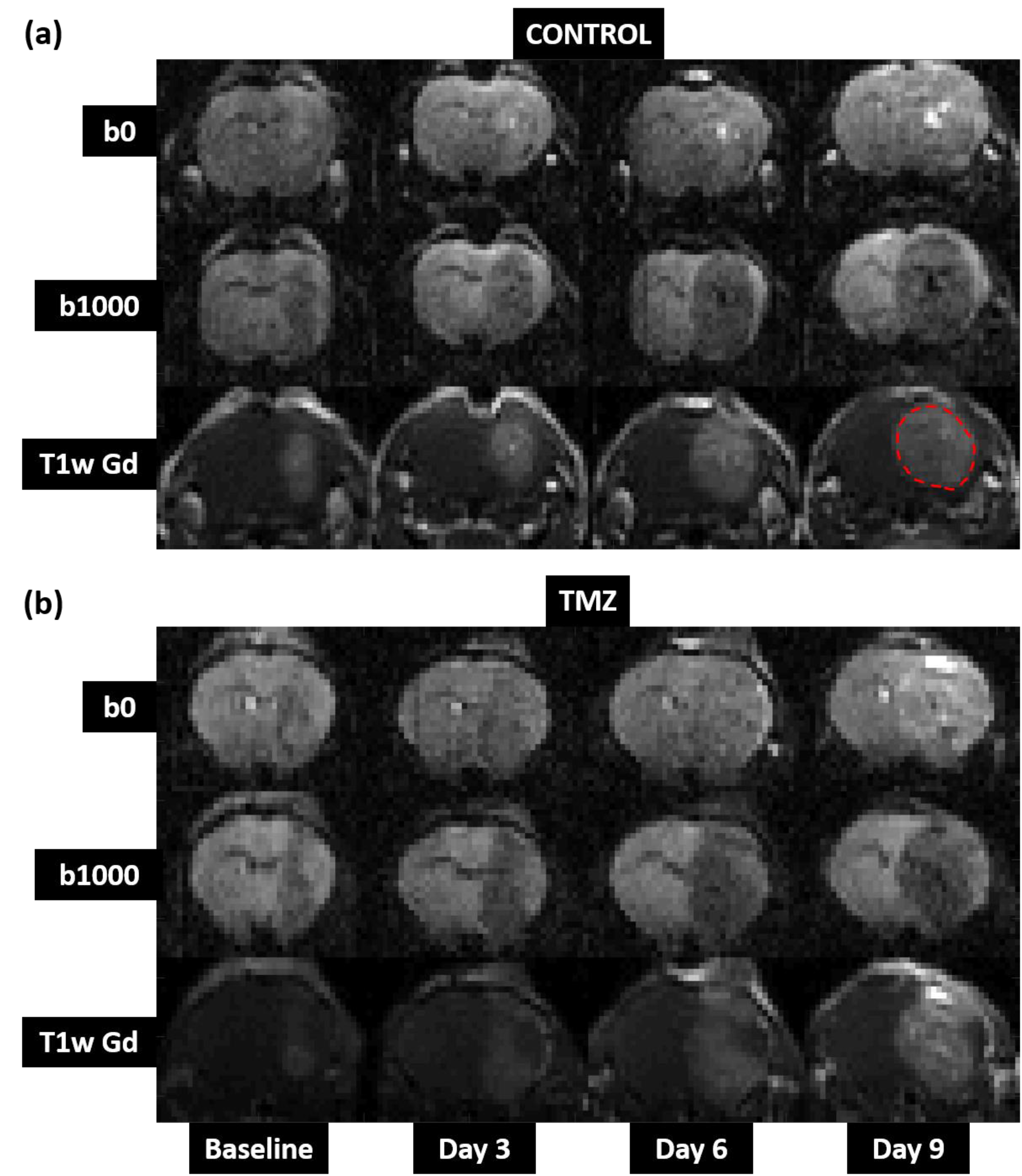
Representative examples of longitudinal structural image data, showing tumour growth from a single mouse in (a) the control cohort and (b) the TMZ-treated cohort. The glioma is isointense with normal brain in T2-weighted images with no diffusion-weighting (b0, top row). Contrast is improved in images with greater diffusion weighting (b1000, middle row), but can still difficult to delineate against normal brain. T1w-gadolinium scans (T1w Gd, bottom row) showed the clearest delineation between tumour and normal brain, and were used to define tumour ROIs (outlined).

Tumour growth increased monotonically with time in both treated and control cohorts (Figure 1). Measurement of tumour volume (based on structural MRI) confirmed this observation (Figure 3a): mean glioma volume was 8 ± 1 mm^3^ at baseline in both cohorts, and after 6-days of treatment, tumour volume more than doubled in both groups of mice (control cohort: 47 ± 2 mm^3^; TMZ-treated cohort: 48 ± 4 mm^3^). Following 9-days of treatment, tumour volumes diverged in the two groups, increasing to 89 ± 7 mm^3^ in the control cohort, compared with 61 ± 8 mm^3^ in TMZ-treated mice. This difference at day 9 was significant (p = 0.029).

### VERDICT quantification of mouse gliomas

Figure 2 shows ADC maps and VERDICT parameter maps, which were spatially heterogeneous. ADC maps of tumours broadly show uniform elevation compared to normal brain, in both the control and TMZ-treated mice. By comparison, VERDICT parameter maps of intracellular volume fraction (*f_ic_*, Figure 2b) and radius (Figure 2e) show clear regions where the parameter values are decreased. Figure S1 shows example, raw diffusion data from a single voxel within a tumour region, and corresponding fits to ADC and VERDICT models.

**Figure 2:**
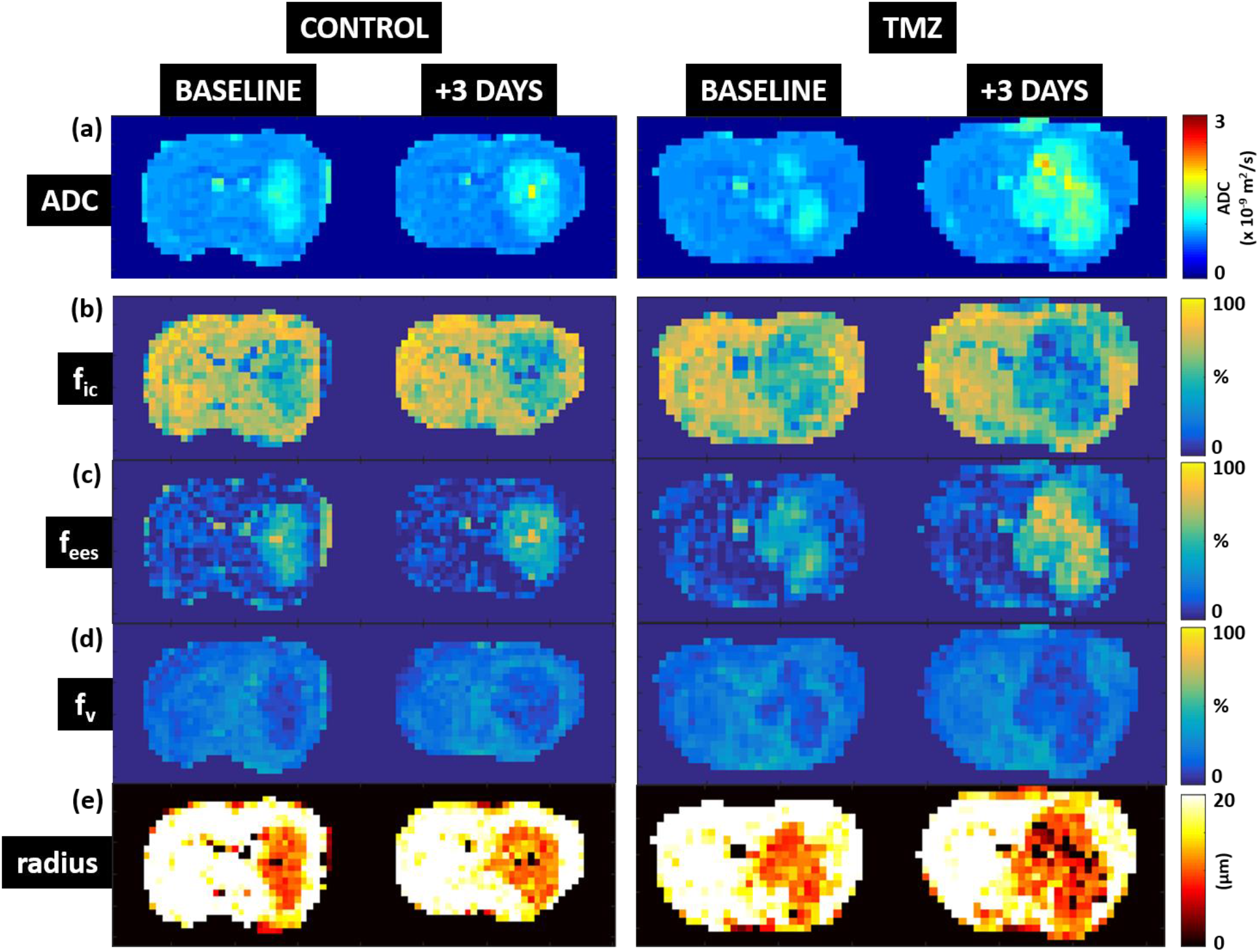
ADC and VERDICT model parameter maps at baseline and 3-days post therapy. (a) apparent diffusion coefficient (ADC), (b) *f_ic_* (intracellular volume fraction), (c) *f_ees_* (extracellular-extravascular volume fraction), (d) *f_v_* (vascular volume fraction), (e) cell radius.

Prior to therapy, VERDICT estimated GL261 tumours to have an intracellular volume fraction (*f_ic_*) of 0.54 ± 0.05 (mean ± SD), extracellular-extravascular volume fraction (*f_ees_*) of 0.39 ± 0.06, vascular volume fraction (*f_v_*) of 0.07 ± 0.02 and radius of 10.6 ± 0.6 μm. Mean ADC prior to therapy was 0.99 ± 0.06 x 10^-9^ m^2^/s.

### Assessment of Temozolomide response with VERDICT

By day 6 of Temozolomide therapy, ADC had significantly increased in the TMZ-treated cohort, compared with the control group (p = 0.001) (Figure 3b). However, VERDICT model parameters reflected changes in microstructure at an earlier time point: control and TMZ-treated cohorts could be distinguished at day 3, based on the cell radius parameter Figure 3f, p = 0.001). The intracellular volume fraction parameter, *f_ic_*, also decreased more rapidly in TMZ-treated tumours than in the control group (Figure 3c, p = 0.005), with a significant difference observed at day 6. An opposite trend was observed in the extracellular volume fraction parameter, *f_ees_*, which steadily increased with tumour growth (Figure 3d, p = 0.005). The vascular volume fraction parameter, *f_v_* (Figure 3c), remained lower than 10% through all timepoints in both cohorts of mice, although at day 6 the parameter was slightly (but significantly) greater in the control group than the TMZ-treated group (p = 0.03).

**Figure 3:**
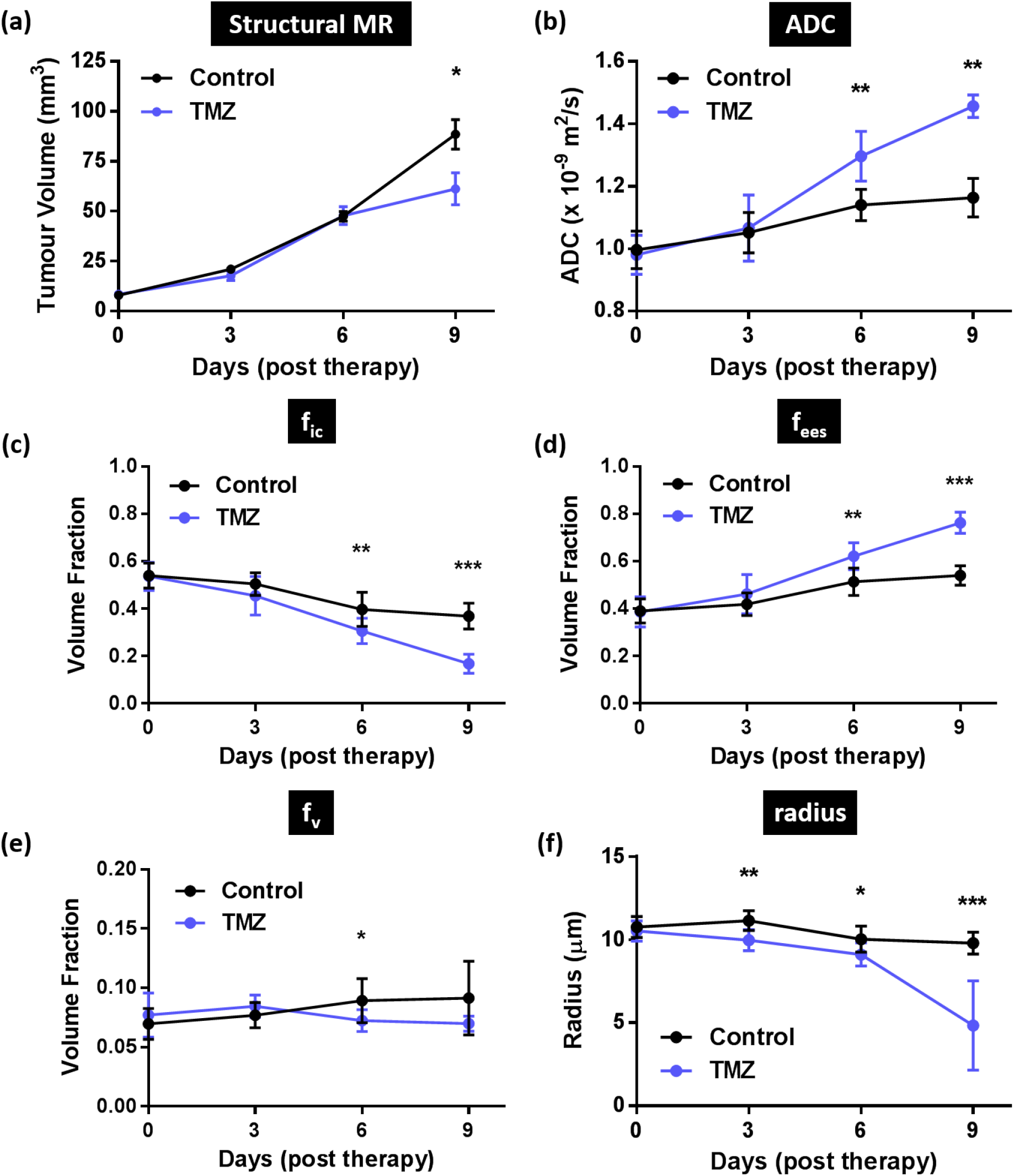
Tumour response to Temozolomide assessed using tumour volume measurements from structural MRI, ADC and VERDICT MRI. (a) Temozolomide had no significant effect on tumour volume until day 9. (b) With ADC, the effect of Temozolomide was observed in the TMZ group at day 6. VERDICT was more sensitive than both measures: a significant difference was observed in the cell radius parameter (f) between the control and TMZ group as early as day 3. A similar divergence between the control and TMZ group was observed in the VERDICT *f_ic_* parameter (c), which decreased more quickly in the TMZ group than in controls.

### VERDICT compared with mouse glioma histology and OPT

After the final imaging timepoint (day 9), histology was performed for comparison with VERDICT parameter maps (Figure 4). Coronal histological sections stained with H&E showed a strong accordance with *f_ic_* and cell radius parameter maps from day 9. In VERDICT maps, tumour regions with a low *f_ic_* value and low cell radius parameter corresponded well with unstained regions of cell death on histology (Figure 4a).

**Figure 4:**
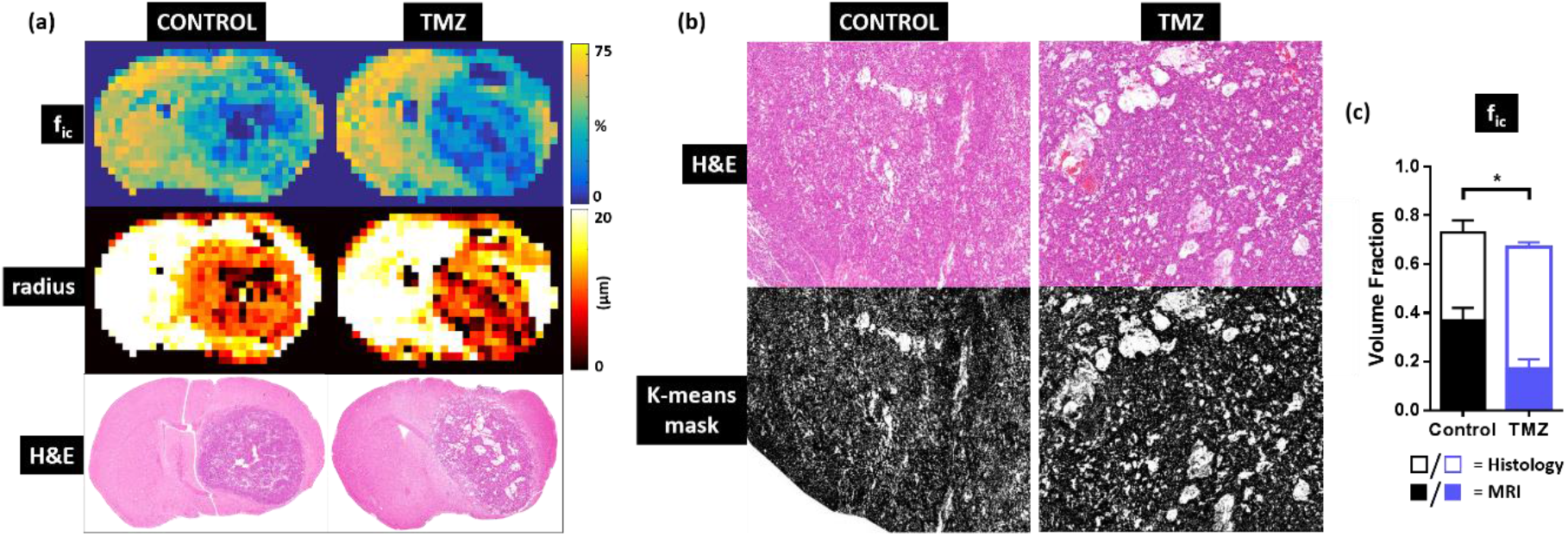
Comparison of coronal sections acquired with VERDICT MRI and histology, and estimation of mouse GBM intracellular volume fraction ( *f_ic_*) from histological sections. (a) Intracellular volume fraction (*f_ic_*) parameter maps, cell radius parameter maps, and H&E stained histology slices. Regions with low *f_ic_* and cell radius match unstained regions on histology. (b) K-means threshold, applied to H&E stained slices. (c) Comparison of *f_ic_* measured in control and TMZ-treated mice, derived from histological analysis and VERDICT MRI.

Quantitative analysis of H&E stained sections was performed to estimate a two-dimensional histological equivalent of the intracellular volume fraction parameter. A k-means threshold was used to mask the stained tissue (black pixels, Figure 4b), from which the ratio of stained tissue to unstained regions was calculated. Histological analysis followed the same trend between control and TMZ-treated animals as VERDICT MRI, where the *f_ic_* parameter was significantly lower in the TMZ-treated group, compared to the control group (Figure 4c, p = 0.02). However, absolute *f_ic_* parameter values from histology (Figure 4c, hollow bars) were more than a factor of 2 greater than *f_ic_* from VERDICT MRI (solid bars).

OPT was also performed after the final imaging time point, on two mouse GBMs labelled with fluorescently-conjugated lectin to estimate a three-dimensional equivalent of the vascular volume fraction parameter, *f_v_*. Blood vessel networks were segmented in a control and a TMZ-treated mouse brain (Figure S2a). The total volume fraction of the blood vessel network was 0.051 ± 0.002 in the control tumour and 0.048 ± 0.001 in the TMZ-treated tumour, which compared well with *f_v_* parameter estimates from VERDICT MRI.

### Characterization of human gliomas with VERDICT

Patient demographics, tumour molecular classification, tumour histology and conventional MRI features are shown in Table S3. The GBMs showed a characteristic tumour mass with an enhancing rim and a non-enhancing necrotic core surrounded by non-enhancing T2-weighted hyperintensity (Figure 5a,b). In comparison, astrocytomas and oligodendrogliomas were more diffuse, with poorly defined boundaries and non-uniform signal intensity on T2-weighted imaging (Figure 5b). ADC was elevated in all tumour types, compared with normal appearing white matter (Figure 5c), and was highest in the tumour core of the GBMs (Figure 6a). The astrocytomas had higher ADC values compared to the oligodendrogliomas.

**Figure 5:**
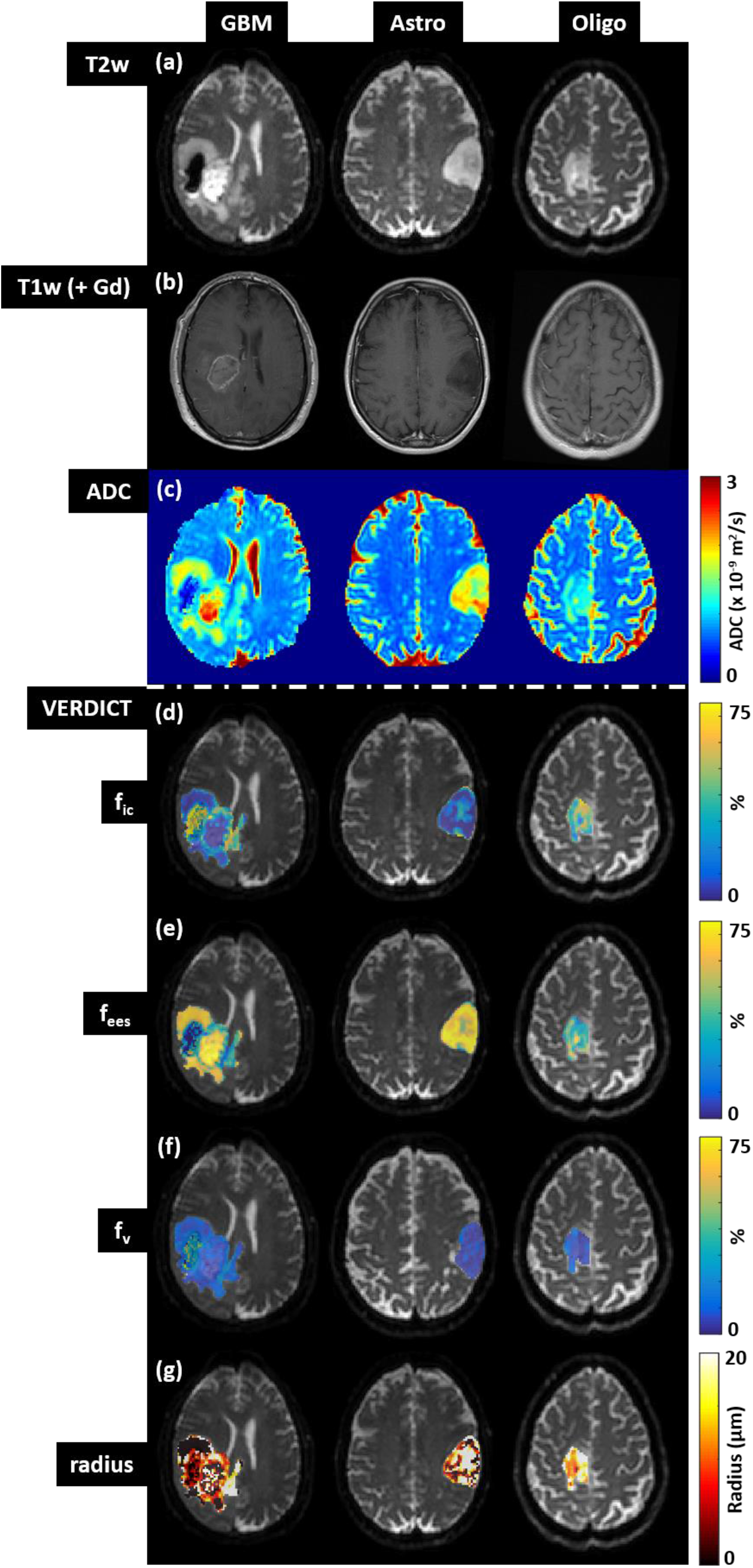
Comparison of (a, b) structural, (c) ADC and (d-g) VERDICT parameter maps in three gliomas: GBM = glioblastoma, Astro = astrocytoma, Oligo = oligodendroglioma.

**Figure 6:**
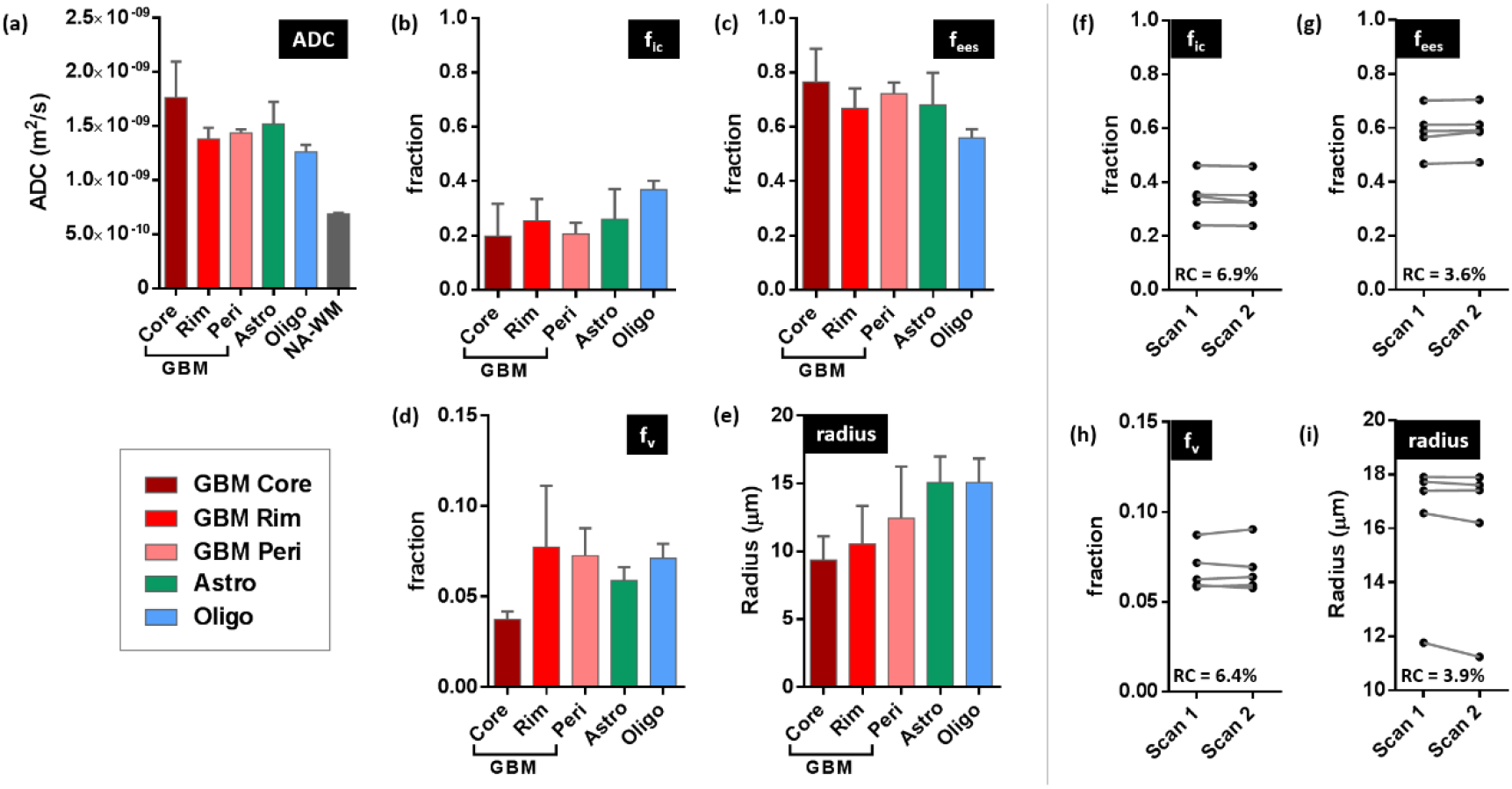
Group analysis (a-e) of ADC and VERDICT parameters in human brain tumours. The four tumour types shown in the key are: GBM = glioblastoma multiforme, Astro = astrocytoma, Oligo = oligodendroglioma. For GBM, analysis was performed on core, rim and peritumoural regions. NA-WM denote normal appearing white matter. (f-i) Repeatability of VERDICT parameters in five brain tumours. No significant difference was measured between trials, in any parameter, indicating a good level of repeatability.

Visual inspection of the GBM VERDICT parameter maps showed a rim with raised intracellular volume fraction (Figure 5d) compared to a central region of relatively low intracellular volume fraction, consistent with a region of higher cell density surrounding a necrotic core. The inverse was observed on extracellular-extravascular volume fraction parameter maps (Figure 5e), where the central tumour core demonstrated elevated extracellular-extravascular space. The tumour rim in GBMs also showed an increased vascular volume fraction compared to the central core (Figure 5f and Figure 6d), consistent with the ring of enhancement evident on T1-weighted post-gadolinium images. The mean cell radius parameters in the tumour core and tumour rim regions of the GBMs (Figure 5e) were consistent with the values measured in the mouse GBM tumour model (~9-10μm, Figure 3f). In the peri-tumour region, this value increased to 12μm.

Interestingly, visual inspection of the VERDICT parameter maps of the peri-tumour regions in the GBMs demonstrated localised regions of heterogeneity. For example, the GBM exhibited a small region of elevated intracellular volume fraction and radius parameter just outside the bulk tumour (yellow arrows, Figure 5d,g), potentially corresponding to regions of invasion.

Visual inspection of the VERDICT volume fraction parameter maps from the astrocytomas and oligodendrogliomas were generally heterogeneous across the full extent of the lesions with regions indicating high and low cell density observed alongside each other. The tumour heterogeneity seen was more apparent on the VERDICT parameter maps than on conventional imaging or the ADC map, suggesting that VERDICT may be more sensitive at reflecting the complex tissue microstructure underlying these tumours. For many voxels in the astrocytomas and oligodendrogliomas, the radius parameter hit the upper boundary of 20μm (white voxels, Figure 5g), indicating that the model fitting for this parameter was unstable in regions of these tumour types. Hence, the mean radius parameters in the astrocytomas and the oligodendrogliomas were considerably higher than for GBMs (Figure 6e).

### Repeatability of VERDICT parameters

The repeatability of the VERDICT parameters was good for all parameters. No significant differences were observed between repeat trials for any of the fitted parameters (Figure 6f-i). The repeatability coefficient was less than 7% for each of the VERDICT parameters, indicating that the fitting was robust and that any variation caused by patient movement was minimised within the image registration pipeline.

### VERDICT compared with patient biopsies

Quantitative analysis of H&E stained biopsies was performed to estimate a nuclei volume fraction for comparison with the VERDICT MRI intracellular volume fraction parameter (Figure S3). For all glioma subtypes, the nuclei volume fraction was lower than *f_ic_* estimated by VERDICT. The oligodendrogliomas had both the largest nuclei volume fraction (Figure S3e) and highest intracellular volume fraction (Figure 5b). There was a positive correlation between the estimates of nuclei volume fraction and VERDICT *f_ic_* parameter (Spearman’s coefficient, r = 0.71, Figure S3f), although it did not reach significance (p = 0.09).

## DISCUSSION

The aim of this study was to evaluate the use of VERDICT MRI to quantify the microstructure of brain tumours, both in a mouse GBM model and in humans, and to assess its ability to detect acute changes induced by chemotherapy (Temozolomide). We have shown that quantitative microstructural parameters from VERDICT MRI reflected changes induced by chemotherapy in mouse brain tumours, earlier than both ADC and standard measurements of tumour volume using structural MRI, and were consistent with measurements from histological sections from the same brains.

Our measurements of tumour microstructure in the GL261 mouse GBM model provided quantitative values, noninvasively, that were consistent with the known structure of brain tumours. At baseline, mean intracellular volume fraction, *f_ic_*, was 0.54 ± 0.05, extracellular-extravascular volume fraction, *f_ees_*, was 0.39 ± 0.06, vascular volume fraction, *f_v_*, was 0.07 ± 0.02 and cell radius was 10.6 ± 0.6 μm.

An increase in tumour volume was observed over time in both cohorts of mice, but was decelerated in those treated with Temozolomide (TMZ). A significant difference in tumour volume, based on structural MRI, was observed between control and TMZ-treated mice at day 9 of the treatment regimen. In both cohorts of mice, ADC increased as the tumours grew, which was most likely due to the onset of necrosis in both treated and control groups, but was even greater in the TMZ-treated group, probably caused by Temzolomide-induced apoptosis. A significant difference in ADC values was observed between the two cohorts of mice after 6-days of treatment.

VERDICT parameters measured in the pre-clinical study also reflected enhanced cell death in the TMZ-treated group, compared to control tumours. Panagiotaki et al. (2014) reported that the induction of apoptosis produced a significant decrease in colorectal carcinoma intracellular cell volume within 6 hours of treatment with gemcitabine (22), and was detectable *in vivo* with VERDICT. In this study, a similar effect was found in brain tumours, although over a longer timescale. Cell radius was the most sensitive of all the VERDICT parameters, producing a significant response to therapy in the TMZ-treated tumours just 3-days after therapy, presumably reflecting a decrease in cell size induced by Temozolomide. This effect was observed three days earlier than any significant changes in ADC and six days earlier than for volumetric measurements. In the control tumours, the mean cell radius parameter was relatively constant throughout the longitudinal study compared to the TMZ-treated tumours, with only a 9.0% decrease in mean radius over the course of the study, compared to a 54.1% decrease in the TMZ-treated mice. Likewise, the intracellular volume fraction decreased more rapidly in the TMZ-treated group compared to the untreated animals, and was significantly lower than controls animals (9.9%) by day 3 of Temozolomide treatment. A much smaller difference of only 1.4% was observed with ADC at the same timepoint. Estimates of vascular volume fraction remained constant throughout the longitudinal study. This suggests that the vasculature was largely unaffected by the Temozolomide, which is in agreement with a previous study by Virrey et al. (29).

Comparison with H&E-stained sections at day-9 post-therapy revealed a strong correspondence between regions of unstained tissue and areas of both low intracellular volume fraction and low cell radius on VERDICT parameter maps, corresponding to regions of greatest necrosis and apoptosis. Quantification of intracellular volume fraction based on H&E slices showed the same trend as VERDICT MRI: cellular density was lower in the TMZ-treated animals compared to controls. However, there was a disparity between the absolute values, which were more than a factor of 2 lower according to VERDICT than from histology. This most likely reflects limitations in the VERDICT model. For example, the model assumes that the cells are impermeable spheres, which is a simplification of the tissue microstructure. Furthermore, the current VERDICT model does not account for the influence of differences in T2 relaxation times between the compartments (30). However, it could also reflect differences between three-dimensional analysis of cell structure (as is inherent within VERDICT), compared to two-dimensional histological analysis.

Following the mouse study, a group of patients were recruited to examine the feasibility of applying VERDICT to human gliomas. Adaptation of the MRI protocol and additional steps in the analysis pipeline were required to translate VERDICT into patients. A reduced diffusion MRI protocol with fewer diffusion shells and directions was implemented to permit a shorter total imaging time of 12 minutes. However, a sufficient number of diffusion weightings and gradient combinations were sampled to maximise coverage at a range of diffusion length scales (7.5-15 μm, based on Einstein’s equation 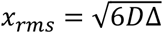, where *D* = 1x10^-9^ and *Δ* = 10-40 ms). Image registration was also required to correct for minor head movement. Finally, voxel-wise fitting was restricted to tumour regions only, firstly, because the pre-clinical study illustrated that VERDICT was an inappropriate model of normal brain parenchyma and, secondly, to reduce image processing time due to the increase in pixel count in patient brains compared to mouse brains. Currently, reliable voxel-wise VERDICT fitting is time consuming over large volumes, but novel algorithms are actively in development to enable VERDICT quantification to be performed faster (31,32).

Based on structural MRI, the tumour phenotype for human GBM was most similar to the GL261 mouse model at day 9. In both mice and patients, a bulk tumour mass was observed containing regions of T2-weighted hyperintensity. However, mouse GBMs enhanced with gadolinium contrast agent across the whole tumour mass, whereas only an area of solid enhancement or rim of enhancement was observed in the GBM patients. There was also no evidence of a peri-tumour region in mice. For ADC and VERDICT parameters, human GBMs were also most similar to GL261 mouse tumours at day 9 compared to the other tumour subtypes. The parameters were actually more comparable with the TMZ-treated cohort of mice rather than the control mice, despite the GBM patients receiving no treatment. Averaged across the equivalent region in GBM patients (tumour core and rim), *f_ic_* was 22% compared to 17% in treated mice; *f_ees_* was 72% compared to 76%; and *f_v_* was 6% compared to 7%. The similarity in these values probably reflects that patients with GBMs tend to present late into development of the tumour, whereas the tumours in the untreated mice at Day 9 have yet to progress to this stage. Moreover, it seems that treatment with Temozolomide has accelerated cell death in the treated cohort to the extent that the volume fractions are comparable with late-stage GBM patients.

VERDICT revealed some interesting sub-regions of heterogeneous tissue microstructure within the peri-tumour zone of the GBMs, which were largely homogenous on the structural images and ADC maps. These regions contained higher *f_ic_*, lower *f_ees_* and higher radius parameter values, indicating an increase in restricted diffusion, typical of increased cellular density. It is possible that this simply reflects a greater mix of neuronal and tumour tissue compared to the bulk of the GBM, however, the lack of uniformity across the peri-tumour region indicates pockets of unusual tissue microstructure. Evidence of localised tumour progression within the peri-tumour region is highly desirable for pre-operative surgical planning (33,34), therefore further imaging with VERDICT in more GBM patients is required to thoroughly interrogate this finding.

The tissue microstructure in the lower grade astrocytomas and oligodendrogliomas was very different to the GBMs. The VERDICT parameters indicated higher cell density compared to GBMs, especially in the oligodendrogliomas: *f_ic_* was increased and *f_ees_* was decreased. The mean radius parameter was considerably higher at 15-16μm, which was caused by many voxels within the astrocytomas and oligodendrogliomas hitting the upper boundary of 20μm. The implementation of VERDICT used here appears to be less appropriate for modelling these tumour subtypes, possibly because of their diffuse phenotype which infiltrates the normal brain parenchyma, which the VERDICT model was not designed to represent. Calcification in oligodendrogliomas is common (and has also been observed in astrocytomas (35)) and known to introduce susceptibility artefacts (36), which may have also affected the diffusion signal. Alternative biophysical models could be investigated and implemented to better approximate these tumour types (37).

Comparison of the VERDICT intracellular volume fraction parameter with biopsy data was also conducted. Intracellular volume fraction could not be estimated as the biopsy samples were highly fragmented. Instead, the cell nuclei volume fraction was compared to the VERDICT intracellular volume fraction parameter, which resulted in a positive correlation as expected. A limitation of this comparison was that the biopsy samples were much smaller than the region of interests drawn on the VERDICT intracellular volume fraction parameter maps and that the exact location of the biopsy region was estimated. This likely explains why the correlation did not reach significance, but even with a relatively small number of patients the positive correlation reaffirms that VERDICT reflects the underlying cell density.

There is a growing literature of evidence suggesting that multi-compartment models of the diffusion MRI signal (22,23,37–39) can provide parameters which more accurately reflect the underlying tissue microstructure and that these models provide higher specificity to pathology compared to more traditional single compartment models such as ADC or DTI. The results of this paper support this consensus. As a potential biomarker of tumour response, VERDICT MRI appears to consistently perform better than structural measurements or ADC. Panagiotaki et al. (2014) also detected decreased intracellular volume fraction in response to gemcitabine chemotherapy (22). However, the clinical part of this study demonstrated that care is required when applying VERDICT to new tumour types. The VERDICT parameters in the patient GBMs appeared to echo the measurements in mouse GBMs, however, the model was less stable when applying the same BallStickSphere to the astrocytomas and oligodendrogliomas. Further studies to optimise VERDICT tissue model for brain tumour subtypes would be invaluable. To this end, ongoing simulations of tumour tissue microenvironments will also be highly important in the optimisation of VERDICT to different tumour types (40).

To summarise, the current standard for assessing brain tumour therapy response using MRI is to make two-dimensional measurements of tumour size based on structural T1- and T2-weighted imaging, as recommended in the RANO Criteria. A host of quantitative techniques for improved measurement of tumour progression are emerging. In this study, we have shown that the VERDICT MRI technique offers superior sensitivity for detecting response to Temozolomide therapy, six days earlier than volumetric alterations are observed and three days earlier than traditional ADC measurements. The intracellular volume fraction, extracellular volume fraction and cell radius parameters estimated by VERDICT all demonstrated changes consistent with cellular apoptosis. We have then adapted and translated this technique into the clinical environment where we imaged a range of glioma subtypes. The microstructural parameters measured in GBM patients were broadly in-line with equivalent parameters in mouse brain GBMs. Evidence of localised increases in cellularity within the peri-tumour region of patient GBMs was observed, however, further work is required to thoroughly characterise the microstructure in these regions. A greater degree of microstructural heterogeneity was observed across the low grade gliomas, reflecting their more diffuse phenotype. Further refinement of the biophysical models is required to robustly characterise these gliomas. In summary, VERDICT is a powerful diffusion MRI technique which offers a host of pathologically-relevant parameters for assessment of brain tumour progression and early treatment response, which can be applied clinically in just over 10 minutes.

## ACKNOWLEDGEMENTS

We acknowledge the support received for the Kings College London & UCL CR-UK and EPSRC Comprehensive Cancer Imaging Centre, in association with the MRC and Department of Health (England) (C1519/A10331) and the Wellcome Trust (WT100247MA); and supported by the National Institute for Health Research University College London Hospitals Biomedical Research Centre; and supported by the Engineering and Physical Sciences Research Council (EPSRC) grant EP/N021967/1.

---

**Table S1:**
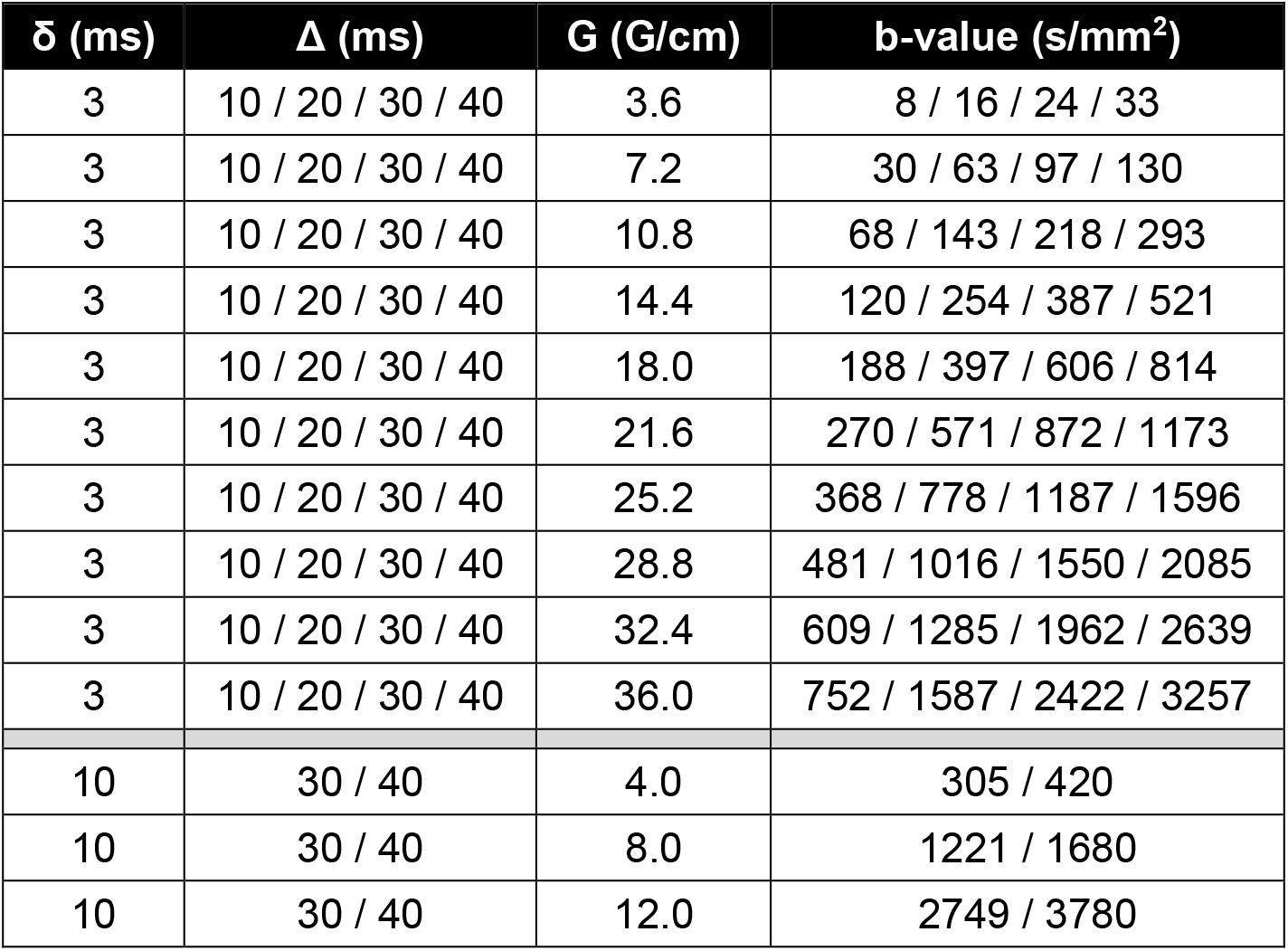
Diffusion gradient combinations used for pre-clinical VERDICT MRI in mouse brains.

| **$\delta$ (ms)** | **$\Delta$ (ms)** | **G (G/cm)** | **b-value (s/mm<sup>2</sup>)** |
| --- | --- | --- | --- |
| 3 | 10 / 20 / 30 / 40 | 3.6 | 8 / 16 / 24 / 33 |
| 3 | 10 / 20 / 30 / 40 | 7.2 | 30 / 63 / 97 / 130 |
| 3 | 10 / 20 / 30 / 40 | 10.8 | 68 / 143 / 218 / 293 |
| 3 | 10 / 20 / 30 / 40 | 14.4 | 120 / 254 / 387 / 521 |
| 3 | 10 / 20 / 30 / 40 | 18.0 | 188 / 397 / 606 / 814 |
| 3 | 10 / 20 / 30 / 40 | 21.6 | 270 / 571 / 872 / 1173 |
| 3 | 10 / 20 / 30 / 40 | 25.2 | 368 / 778 / 1187 / 1596 |
| 3 | 10 / 20 / 30 / 40 | 28.8 | 481 / 1016 / 1550 / 2085 |
| 3 | 10 / 20 / 30 / 40 | 32.4 | 609 / 1285 / 1962 / 2639 |
| 3 | 10 / 20 / 30 / 40 | 36.0 | 752 / 1587 / 2422 / 3257 |
| 10 | 30 / 40 | 4.0 | 305 / 420 |
| 10 | 30 / 40 | 8.0 | 1221 / 1680 |
| 10 | 30 / 40 | 12.0 | 2749 / 3780 |

**Table S2:** Diffusion gradient combinations used for VERDICT MRI in patients.

| **$\delta$ (ms)** | **$\Delta$ (ms)** | **G (G/cm)** | **b-value (s/mm<sup>2</sup>)** |
| --- | --- | --- | --- |
| 3.2 | 15.6 | 8.7 | 80 |
| 4.3 | 16.6 | 8.9 | 160 |
| 6.0 | 18.3 | 9.1 | 350 |
| 7.5 | 19.9 | 9.3 | 600 |
| 17.8 | 28.0 | 4.5 | 1000 |
| 20.9 | 31.1 | 4.5 | 1500 |
| 12.6 | 24.9 | 9.2 | 2000 |
| 13.8 | 26.1 | 9.2 | 2500 |
| 16.8 | 44.4 | 6.2 | 3000 |

**Table S3:** Summary of patient brain tumour histopathology and MRI characteristics.

| Patient | Sex | Age | Histopathology | Genetics | Location | MRI |
| --- | --- | --- | --- | --- | --- | --- |
| 1 | M | 52 | Glioblastoma<br>WHO IV | IDH mutation<br>1p/19q<br>uncodeleted | L insula | Solid enhancement with<br>surrounding T2W<br>hyperintensity.<br>Restricted diffusion of<br>enhancing component. |
| 2 | F | 68 | Glioblastoma<br>WHO IV | IDH<br>unmutated<br>1p/19q<br>uncodeleted | L frontal | Rim enhancing lesion<br>with central necrosis and<br>surrounding T2W<br>hyperintensity.<br>Restricted diffusion of the<br>enhancing rim.<br>Areas of intralesional<br>haemorrhage. |
| 3 | M | 60 | Glioblastoma<br>WHO IV | IDH<br>Unmutated<br>1p/19q<br>uncodeleted | R fronto-<br>parietal | Rim enhancing lesion<br>with central necrosis and<br>surrounding T2W<br>hyperintensity.<br>Restricted diffusion of the<br>enhancing rim.<br>Areas of intralesional<br>haemorrhage and large<br>haematoma. |
| 4 | F | 50 | Anaplastic<br>astrocytoma<br>WHO III | IDH mutation<br>1p/19q<br>uncodeleted | L frontal | T2W hyperintense.<br>Patchy enhancement.<br>Mixed diffusion. |
| 5 | M | 37 | Diffuse astrocytoma<br>WHO II | IDH mutation<br>1p/19q<br>uncodeleted | L insula | T2W hyperintense.<br>Non-enhancing.<br>Facilitated diffusion. |
| 6 | M | 43 | Oligodendroglioma<br>WHO II | IDH mutation<br>1p/19q<br>codeleted | R frontal | T2W hyperintense.<br>Patchy enhancement.<br>Mixed diffusion. |
| 7 | M | 22 | Oligodendroglioma<br>WHO II | IDH mutated<br>1p/19q<br>codeleted | R frontal | T2W hyperintense.<br>Predominantly non-enhancing.<br>Facilitated diffusion |
| 8 | F | 52 | Oligodendroglioma<br>WHO II | IDH mutation<br>1p/19q<br>codeleted | L fronto-<br>temporal | T2W hyperintense.<br>Patchy enhancement.<br>Predominantly facilitated<br>diffusion.<br>Areas of calcification. |
| 9 | M | 35 | Clinical assessment<br>only – Astrocytoma | - | L frontal | T2W hyperintense.<br>Patchy enhancement.<br>Mixed diffusion. |

**Figure S1:**
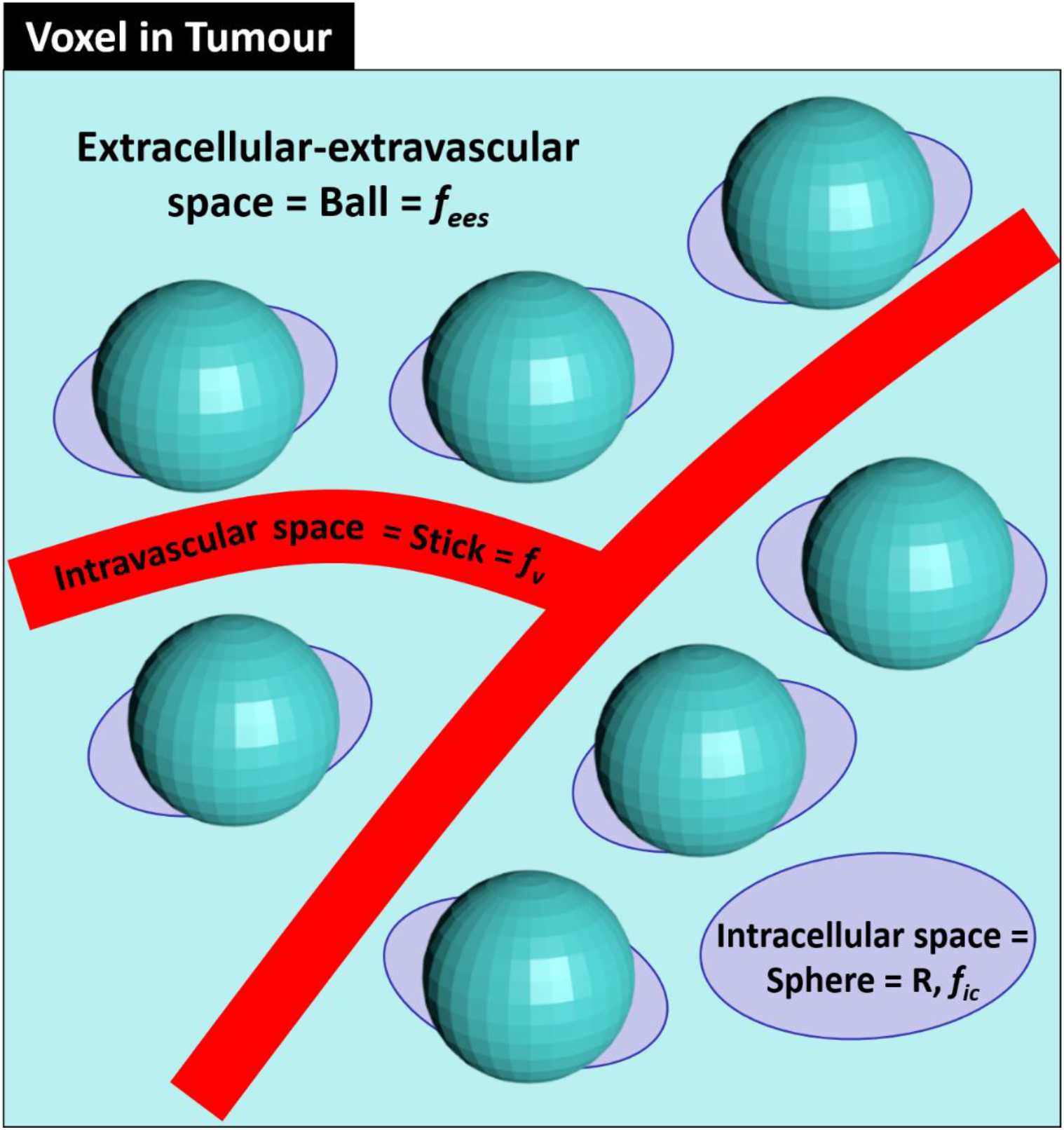
Schematic showing how the VERDICT BallSphereStick model represents the microstructure within a voxel of tumour tissue. The extracellular-extravascular space (*f_ees_*) is represented as a Ball compartment, the intracellular space (*f_ic_*) is represented as a Sphere compartment (with average radius *R*) and the intravascular space (*f_v_*) is represented as a Stick. Further details on the functional form of each component can be found in Panagiotaki et al (1).

**Figure S2:**
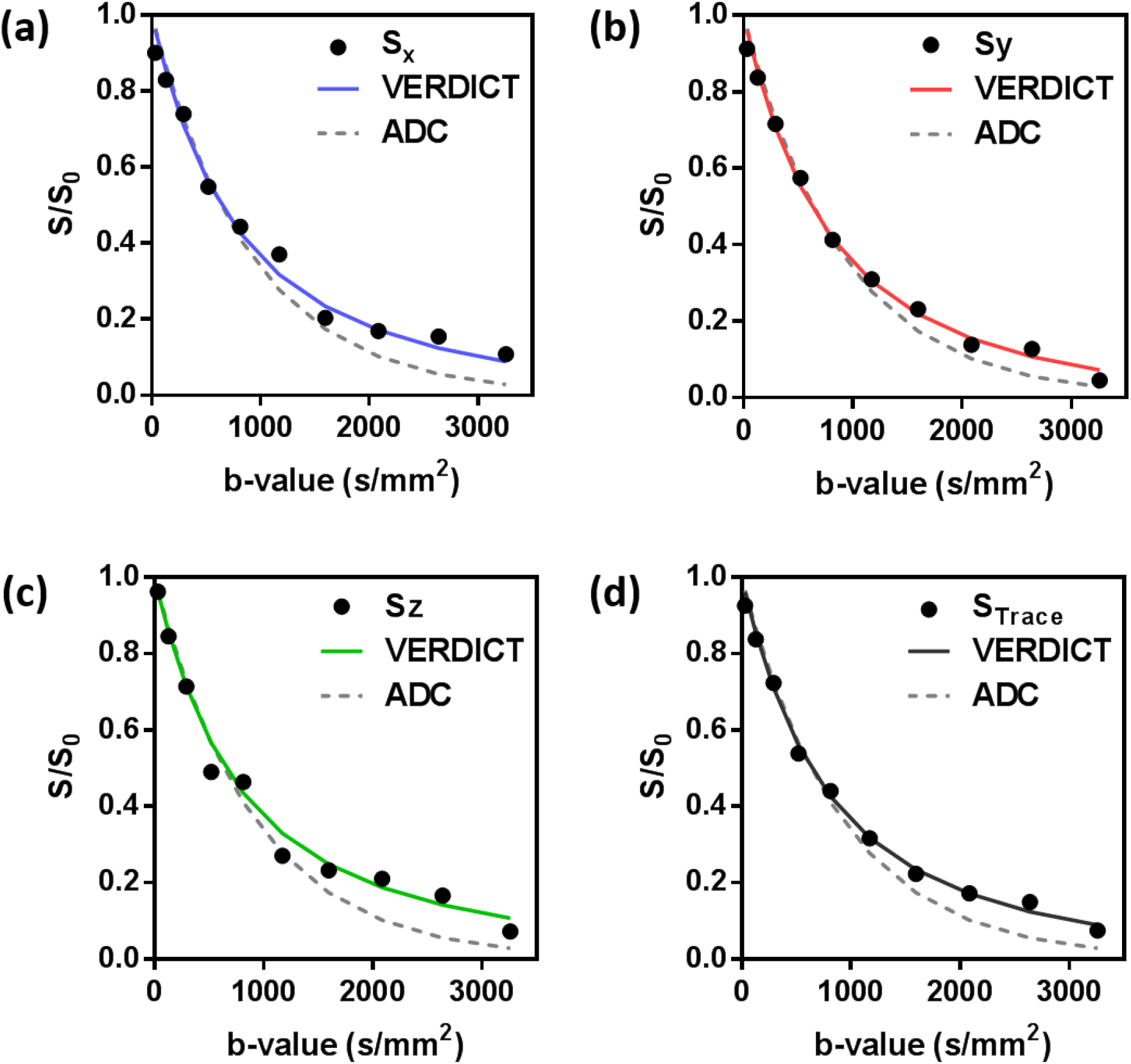
Examples of DWI data fitted to the VERDICT model (solid lines) and ADC model (dashed lines), taken from a single voxel within a GL261 mouse brain tumour ROI. The individual graphs show data acquired with diffusion gradients orientated in the (a) x-direction, (b) y-direction, (c) z-direction. (d) shows the trace diffusion signal, with its corresponding fits to both models.

**Figure S3:**
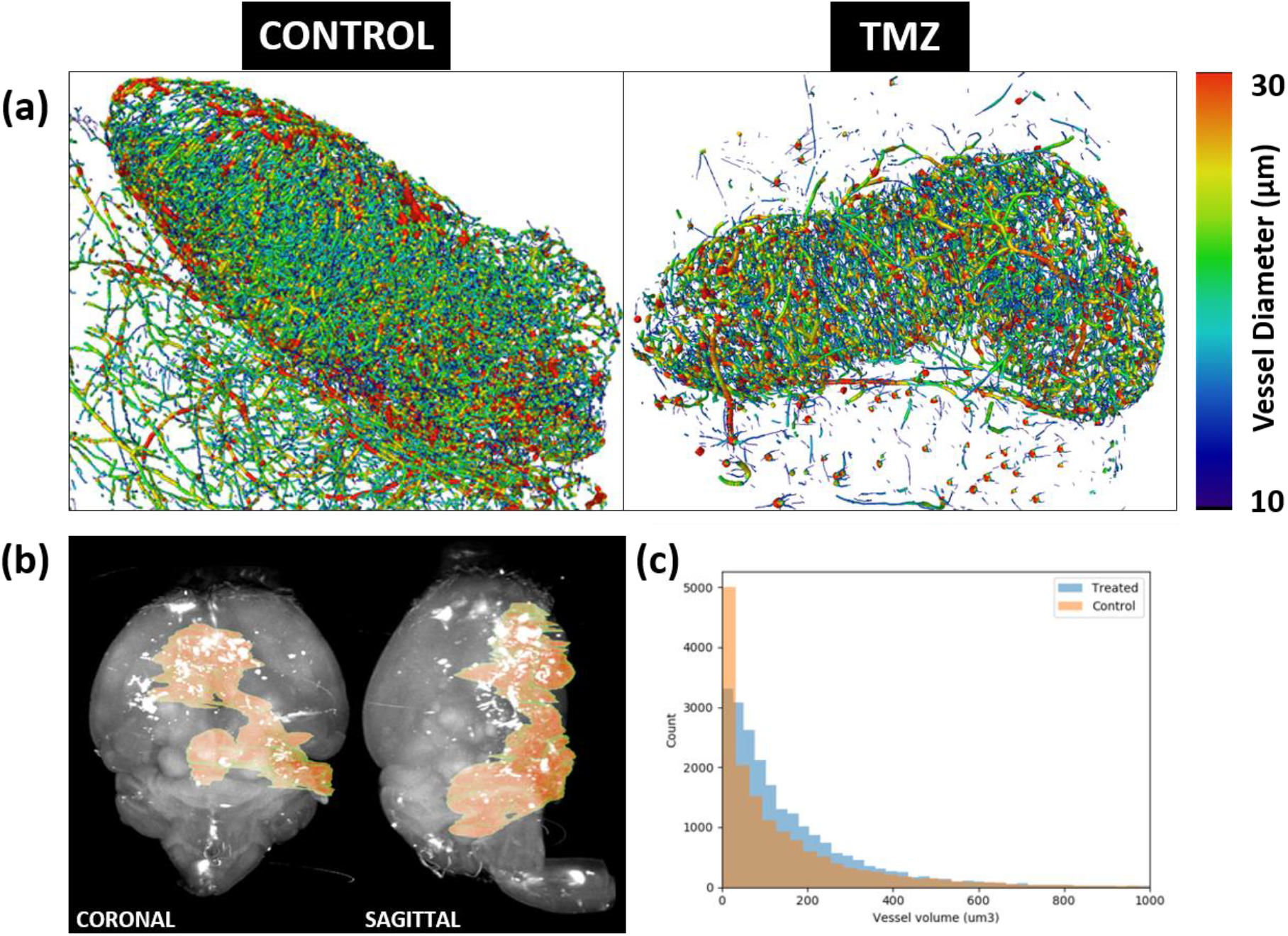
Estimation of mouse GBM (GL261) vascular volume fraction ( *f_v_*) from optical projection tomography data. (a) Blood vessel networks from a control and TMZ-treated mouse brain tumour. (b) OPT volume render from a control mouse brain showing the extent of tumour (orange). (c) Comparison of blood vessel volume histograms from a control and a TMZ-treated mouse GBM.

**Figure S4:**
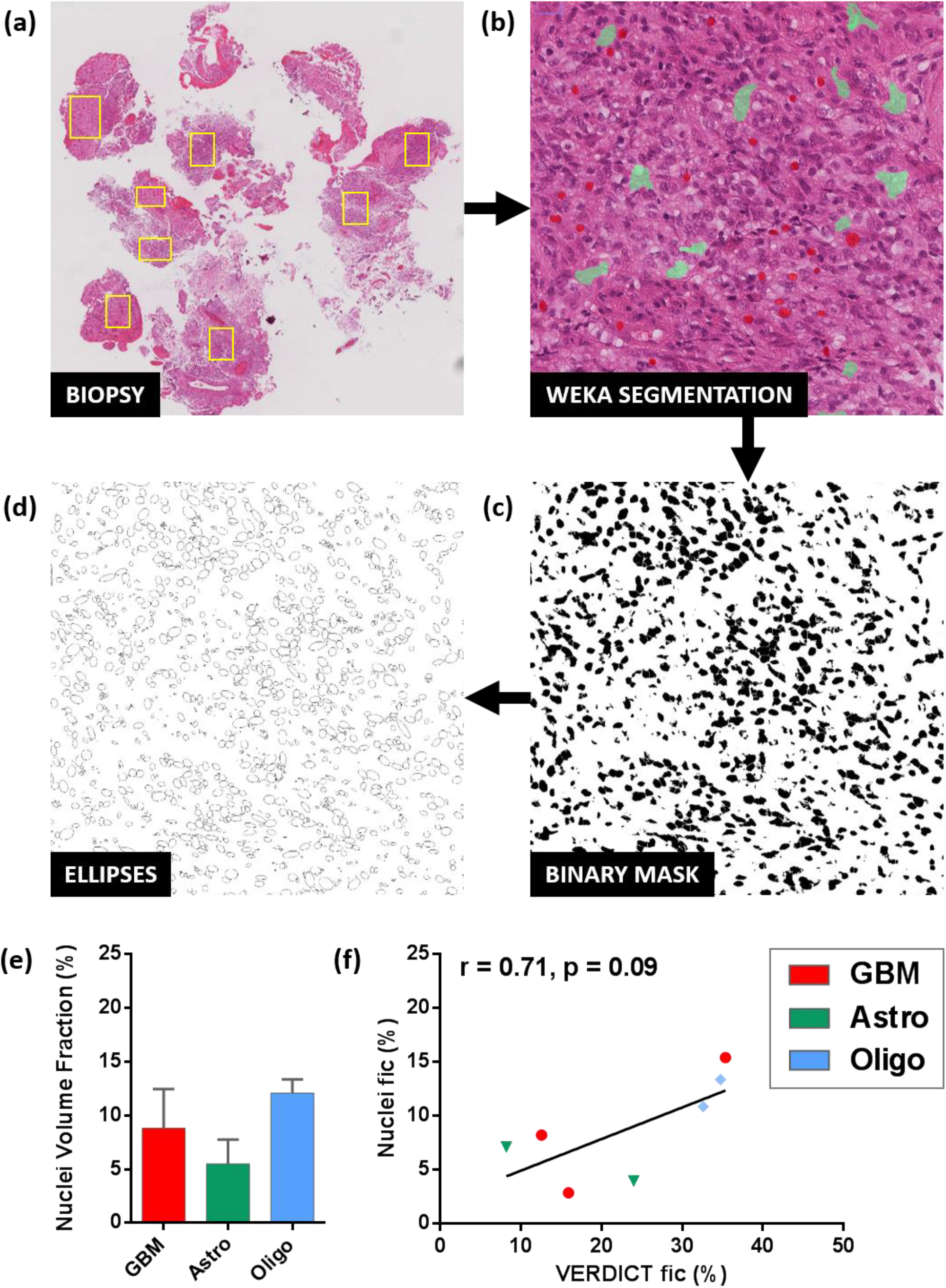
Patient biopsy nuclei volume fraction analysis pipeline and results. (a) Tumour regions (yellow outlines) within biopsies were sampled and saved at 40x magnification (0.25μm/pixel). (b) Manual classification of biopsy samples into nuclei (red) and ‘everything else’ (green) (c) Binary mask of nuclei produced from the Weka Segmentation machine learning tool. (d) Ellipses generated from the binary mask. (e) Mean cell nucleus volume fraction estimated from biopsy samples. (f) Linear correlation of VERDICT intracellular volume fraction parameter (*f_ic_*) against cell nucleus volume fraction from biopsy samples (r = Spearman’s coefficient).

## SUPPLEMENTARY MATERIALS AND METHODS

### Optical Projection Tomography – Preparation, Imaging and Image Processing

Immediately after the final MRI scan, mice were intravenously-injected (via the tail vein) with 100 μg lectin-AlexFluor 647 (Thermo Fisher Scientific, L32451), diluted in sterile saline at neutral pH (100 μl) containing 1 mM CaCl_2_ in order to fluorescently label blood vessels, which was allowed to circulate for 5 minutes. Mice were then terminally anaesthetised via intraperitoneal injection of 100 mg/kg sodium pentobarbital (Pentoject, Animalcare, York, UK), diluted in 0.1 ml phosphate buffered saline (PBS).

Next, heparin (Wockhardt, Heparin Sodium) diluted in saline was administered by intraperitoneal injection (0.2 ml, with 1000 IU/ml). Mice were perfuse-fixed by opening along the sternum to expose the heart. A small puncture was made in the apex of the heart using micro scissors, and a blunted 25G butterfly cannula was inserted through the left ventricle and into the ascending aorta, where it was secured with a ligature. PBS (30 ml, maintained at 37 °C) was perfused into the heart at 3 ml/min with a perfusion pump (Watson Marlow, 5058). After complete drainage of blood, 40 ml of 4% paraformaldehyde (PFA, VWR chemicals) was perfused.

Intact mouse brains were extracted from the skull and stored for 12 hours in 4% PFA, at 4 °C. Following perfuse-fixation, tumours were rinsed three times in PBS, for 10 minutes each, to remove residual formaldehyde (2). Brain tissue was then optically cleared with BABB (1:2 Benzyl alcohol: Benzyl benzoate), by immersion in methanol for 48 hours followed by immersion in BABB for 48 hours.

Fluorescence within vasculature was visualized with OPT (Bioptonics, MRC Technologies, Edinburgh) using an exposure time of 1600-2000 ms and a rotation step of 0.45 degrees (Figure S2b). The final x-y resolution ranged from 4.3 μm to 8 μm, depending on sample size (3). Data were reconstructed using NRecon software (SkyScan, Kontich, Belgium).

Brains were segmented from background by simple thresholding. Tumour tissue was identified by manually drawing a three-dimensional region of interest (ROI) (Amira, FEI, Oregon, US). Background autofluorescence was removed from OPT images by applying a 3-dimensional Gaussian filter of width 25 pixels. Filtered data were subtracted from an unfiltered copy of the data. A Frangi filter was applied to enhance vessel structure. Finally, the data were thresholded to segment vascular structures from background. Data were visualised in three-dimensions using Amira (Figure S2a). Blood volume was estimated as the ratio of total vascular volume to tumour ROI volume.

### Patient Biopsy Image Processing

Biopsies were retrieved from seven of the brain tumour patients. Image processing was performed on H&E-stained samples to estimate nuclei volume fraction for comparison with VERDICT estimates of intracellular volume fraction. Figure S3 shows the image processing pipeline used to extract nuclei from the images. First, tumour regions within the biopsy slides were sampled and saved as TIFF files at 40x magnification (Figure S3a), typically containing 5000 pixels^2^ (0.25μm/pixel). Using the Weka Segmentation plug-in (4) for Fiji (NIH, US) (5), two classifiers were trained to represent cell nuclei (Figure S3b, red) and ‘everything else’ (green). The same classifier was used for all patient biopsy samples. A binary mask was generated for each biopsy sample (Figure S3c), which was then converted to ellipses (Figure S3d) using the in-built function within Fiji. The ellipses were constrained to have an area, A > 5um^2^ and A < 200um^2^ to eliminate noise and any abnormally large ellipses. Nuclei volume fraction was estimated as the ratio of total volume across all ellipses to total volume of the original image.

## Notes

**Conflict of Interest Statement**: The authors declare no potential conflicts of interest.

## REFERENCES

1. Stupp R, Hegi ME, Mason WP, van den Bent MJ, Taphoorn MJ, Janzer RC, Ludwin SK, Allgeier A, Fisher B, Belanger K. Effects of radiotherapy with concomitant and adjuvant temozolomide versus radiotherapy alone on survival in glioblastoma in a randomised phase III study: 5-year analysis of the EORTC-NCIC trial. The lancet oncology 2009;10(5):459–466.

2. Desouza R, Shaweis H, Han C, Sivasubramiam V, Brazil L, Beaney R, Sadler G, Alsarraj S, Hampton T, Logan J. Has the survival of patients with glioblastoma changed over the years? The British Journal of Cancer 2016;114(2):146.

3. Hottinger AF, Stupp R, Homicsko K. Standards of care and novel approaches in the management of glioblastoma multiforme. Chinese journal of cancer 2014;33(1):32.

4. Macdonald DR, Cascino TL, Schold SC Jr, Cairncross JG. Response criteria for phase II studies of supratentorial malignant glioma. Journal of Clinical Oncology 1990;8(7):1277–1280.

5. Young R, Gupta A, Shah A, Graber J, Zhang Z, Shi W, Holodny A, Omuro A. Potential utility of conventional MRI signs in diagnosing pseudoprogression in glioblastoma. Neurology 2011;76(22):1918–1924.

6. Verma N, Cowperthwaite MC, Burnett MG, Markey MK. Differentiating tumor recurrence from treatment necrosis: a review of neuro-oncologic imaging strategies. Neuro-oncology 2013;15(5):515–534.

7. Norden AD, Drappatz J, Wen PY. Novel anti-angiogenic therapies for malignant gliomas. The Lancet Neurology 2008;7(12):1152–1160.

8. Narayana A, Kelly P, Golfinos J, Parker E, Johnson G, Knopp E, Zagzag D, Fischer I, Raza S, Medabalmi P. Antiangiogenic therapy using bevacizumab in recurrent high-grade glioma: impact on local control and patient survival. Journal of neurosurgery 2009;110(1):173–180.

9. Wen PY, Macdonald DR, Reardon DA, Cloughesy TF, Sorensen AG, Galanis E, DeGroot J, Wick W, Gilbert MR, Lassman AB. Updated response assessment criteria for high-grade gliomas: response assessment in neuro-oncology working group. Journal of Clinical Oncology 2010;28(11):1963–1972.

10. Metellus P, Coulibaly B, Colin C, de Paula AM, Vasiljevic A, Taieb D, Barlier A, Boisselier B, Mokhtari K, Wang XW. Absence of IDH mutation identifies a novel radiologic and molecular subtype of WHO grade II gliomas with dismal prognosis. Acta neuropathologica 2010;120(6):719–729.

11. Maier SE, Sun Y, Mulkern RV. Diffusion imaging of brain tumors. Nmr Biomed 2010;23(7):849–864.

12. Sugahara T, Korogi Y, Kochi M, Ikushima I, Shigematu Y, Hirai T, Okuda T, Liang L, Ge Y, Komohara Y. Usefulness of diffusion‐weighted MRI with echo‐planar technique in the evaluation of cellularity in gliomas. Journal of Magnetic Resonance Imaging 1999;9(1):53–60.

13. Murakami R, Hirai T, Sugahara T, Fukuoka H, Toya R, Nishimura S, Kitajima M, Okuda T, Nakamura H, Oya N. Grading astrocytic tumors by using apparent diffusion coefficient parameters: superiority of a one-versus two-parameter pilot method. Radiology 2009;251(3):838–845.

14. LaViolette PS, Mickevicius NJ, Cochran EJ, Rand SD, Connelly J, Bovi JA, Malkin MG, Mueller WM, Schmainda KM. Precise ex vivo histological validation of heightened cellularity and diffusion-restricted necrosis in regions of dark apparent diffusion coefficient in 7 cases of high-grade glioma. Neuro-oncology 2014;16(12):1599–1606.

15. Hein PA, Eskey CJ, Dunn JF, Hug EB. Diffusion-weighted imaging in the follow-up of treated high-grade gliomas: tumor recurrence versus radiation injury. American Journal of Neuroradiology 2004;25(2):201–209.

16. Lee WJ, Choi SH, Park C-K, Yi KS, Kim TM, Lee S-H, Kim J-H, Sohn C-H, Park S-H, Kim IH. Diffusion-weighted MR imaging for the differentiation of true progression from pseudoprogression following concomitant radiotherapy with temozolomide in patients with newly diagnosed high-grade gliomas. Academic radiology 2012;19(11):1353–1361.

17. Carr HY, Purcell EM. Effects of diffusion on free precession in nuclear magnetic resonance experiments. Physical review 1954;94(3):630.

18. Le Bihan D, Breton E, Lallemand D, Grenier P, Cabanis E, Laval-Jeantet M. MR imaging of intravoxel incoherent motions: application to diffusion and perfusion in neurologic disorders. Radiology 1986;161(2):401–407.

19. Basser PJ, Mattiello J, LeBihan D. MR diffusion tensor spectroscopy and imaging. Biophysical journal 1994;66(1):259–267.

20. Lu H, Jensen JH, Ramani A, Helpern JA. Three‐dimensional characterization of non‐gaussian water diffusion in humans using diffusion kurtosis imaging. Nmr Biomed 2006;19(2):236–247.

21. Zhang H, Schneider T, Wheeler-Kingshott CA, Alexander DC. NODDI: practical in vivo neurite orientation dispersion and density imaging of the human brain. Neuroimage 2012;61(4):1000–1016.

22. Panagiotaki E, Walker-Samuel S, Siow B, Johnson SP, Rajkumar V, Pedley RB, Lythgoe MF, Alexander DC. Noninvasive quantification of solid tumor microstructure using VERDICT MRI. Cancer research 2014;74(7):1902–1912.

23. Panagiotaki E, Chan RW, Dikaios N, Ahmed HU, O’callaghan J, Freeman A, Atkinson D, Punwani S, Hawkes DJ, Alexander DC. Microstructural characterization of normal and malignant human prostate tissue with vascular, extracellular, and restricted diffusion for cytometry in tumours magnetic resonance imaging. Investigative radiology 2015;50(4):218–227.

24. Szatmári T, Lumniczky K, Désaknai S, Trajcevski S, Hídvégi EJ, Hamada H, Sáfrány G. Detailed characterization of the mouse glioma 261 tumor model for experimental glioblastoma therapy. Cancer science 2006;97(6):546–553.

25. Workman P, Aboagye EO, Balkwill F, Balmain A, Bruder G, Chaplin DJ, Double JA, Everitt J, Farningham DAH, Glennie MJ, Kelland LR, Robinson V, Stratford IJ, Tozer GM, Watson S, Wedge SR, Eccles SA, Navaratnam V, Ryder S, Inst NCR. Guidelines for the welfare and use of animals in cancer research. Brit J Cancer 2010;102(11):1555–1577.

26. Cook P, Bai Y, Nedjati-Gilani S, Seunarine K, Hall M, Parker G, Alexander D. Camino: open-source diffusion-MRI reconstruction and processing.

27. Mohammadi S, Möller HE, Kugel H, Müller DK, Deppe M. Correcting eddy current and motion effects by affine whole‐brain registrations: Evaluation of three‐dimensional distortions and comparison with slicewise correction. Magnetic Resonance in Medicine 2010;64(4):1047–1056.

28. Penny WD, Friston KJ, Ashburner JT, Kiebel SJ, Nichols TE. Statistical parametric mapping: the analysis of functional brain images: Academic press; 2011.

29. Virrey JJ, Golden EB, Sivakumar W, Wang W, Pen L, Schönthal AH, Hofman FM, Chen TC. Glioma-associated endothelial cells are chemoresistant to temozolomide. Journal of neuro-oncology 2009;95(1):13–22.

30. Fournet G, Li J-R, Le Bihan D, Ciobanu L. The influence of acquisition parameters on the metrics of the bi-exponential IVIM model. 2016.

31. Bonet-Carne E, Daducci A, Panagiotaki E, Johnston E, Stevens N, Atkinson D, Punwani S, Alexander DC. Non-invasive quantification of prostate cancer using AMICO framework for VERDICT MRI. 2016. International Society for Magnetic Resonance in Medicine (ISMRM).

32. Harms R, Fritz F, Tobisch A, Goebel R, Roebroeck A. Robust and fast nonlinear optimization of diffusion MRI microstructure models. NeuroImage 2017;155:82–96.

33. Price S, Gillard J. Imaging biomarkers of brain tumour margin and tumour invasion. The British journal of radiology 2011;84(special_issue_2):S159–S167.

34. Ahmed R, Oborski MJ, Hwang M, Lieberman FS, Mountz JM. Malignant gliomas: current perspectives in diagnosis, treatment, and early response assessment using advanced quantitative imaging methods. Cancer management and research 2014;6:149.

35. Makariou E, Patsalides AD. Intracranial calcifications. Appl Radiol 2009;38(11):48–50.

36. Smits M. Imaging of oligodendroglioma. The British journal of radiology 2016;89(1060):20150857.

37. Panagiotaki E, Schneider T, Siow B, Hall MG, Lythgoe MF, Alexander DC. Compartment models of the diffusion MR signal in brain white matter: a taxonomy and comparison. Neuroimage 2012;59(3):2241–2254.

38. Johnston E, Pye H, Bonet-Carne E, Panagiotaki E, Patel D, Galazi M, Heavey S, Carmona L, Freeman A, Trevisan G. INNOVATE: A prospective cohort study combining serum and urinary biomarkers with novel diffusion-weighted magnetic resonance imaging for the prediction and characterization of prostate cancer. BMC cancer 2016;16(1):816.

39. Colgan N, Siow B, O’callaghan J, Harrison I, Wells JA, Holmes H, Ismail O, Richardson S, Alexander DC, Collins E. Application of neurite orientation dispersion and density imaging (NODDI) to a tau pathology model of Alzheimer’s disease. NeuroImage 2016;125:739–744.

40. Ben Hipwell TR, Paul Sweeney, Angela D’Esposito, Morium Ali, Eleftheria Panagiotaki, Mark Lythgoe, Daniel Alexander, Rebecca Shipley, Simon Walker-Samuel. A novel framework for simulating the in-vivo diffusion MRI signal in solid tumours, based on high-resolution optical imaging data from real-world tumours. 2017.

## References

1. Panagiotaki E, Schneider T, Siow B, Hall MG, Lythgoe MF, Alexander DC. Compartment models of the diffusion MR signal in brain white matter: a taxonomy and comparison. Neuroimage 2012;59(3):2241–2254.

2. Janssen FJ. A study of the absorption and scattering factors of light in whole blood. Medical & biological engineering 1972;10(2):231–240.

3. Jonkman JE, Swoger J, Kress H, Rohrbach A, Stelzer EH. Resolution in optical microscopy. Methods in enzymology 2003;360:416–446.

4. Arganda-Carreras I, Kaynig V, Rueden C, Eliceiri KW, Schindelin J, Cardona A, Sebastian Seung H. Trainable Weka Segmentation: a machine learning tool for microscopy pixel classification. Bioinformatics 2017:btx180.

5. Schindelin J, Arganda-Carreras I, Frise E, Kaynig V, Longair M, Pietzsch T, Preibisch S, Rueden C, Saalfeld S, Schmid B. Fiji: an open-source platform for biological-image analysis. Nature methods 2012;9(7):676–682.

